# Early gonadotoxic effects of cyclophosphamide on the prepubertal testis and the feasibility of reducing toxicity through combined antioxidant therapy

**DOI:** 10.1101/2025.03.18.643799

**Authors:** Amirhesam Eskafinoghani, Arturo Reyes Palomares, Xia Hao, Roudabeh Mohammadi, Arian Lundberg, Kenny A. Rodriguez-Walberg

**Affiliations:** Department of Oncology-Pathology, Karolinska Institute, Stockholm, Sweden; Laboratory of Translational Fertility Preservation, Karolinska Institute, Stockholm, Sweden; Departamento de especialidades Quirúrgicas, Bioquímica e Inmunología, Facultad de Medicina, Universidad de Málaga, Malaga, Spain; Department of Protein Science, SciLifeLab, KTH Royal Institute of Technology, Stockholm, Sweden; Department of Reproductive Medicine, Division of Gynecology and Reproduction, Karolinska University Hospital, Stockholm, Sweden

## Abstract

**One Sentence Summary:** Early gonadotoxic effects of cyclophosphamide (CPA) on prepubertal testes were identified, with molecular damage detectable within 16 hours and histological changes by 32 hours, while antioxidant co-treatment showed potential for protection. CPA induces early molecular damage in prepubertal testes, with protective antioxidant effects observed within the first 16 hours, highlighting the potential for mitigating testicular toxicity without compromising anticancer efficacy. This study demonstrates that CPA-induced testicular toxicity in prepubertal mice occurs early at the transcriptional level, with antioxidants showing protective effects, underscoring the need for further research on safe and effective protective strategies.

At high doses, alkylating drugs’ toxicity leads to testicular atrophy and spermatogonial stem cells depletion, rendering survivors of childhood cancers infertile, as a recognized late effect. We used in-vivo experimental models to identify early gonadotoxic effects induced by cyclophosphamide (CPA) on prepubertal testes and to investigate if a treatment with antioxidants could reduce that toxicity. Effects were estimated by histological evaluation and whole-transcriptome RNA-sequencing (RNA-seq) of sacrificed animals at 8 hours intervals. Sixty pre-pubertal mice were randomly assigned to 1) intraperitoneal treatment with 100 mg/Kg CPA; 2) CPA in addition to antioxidants (AO) L-Carnitine (LC 50mg/kg) and N-acetyl cysteine (NAC 50); 3) AO alone; 4) control receiving saline solution (Ctrl) (N=15 for each group). Three additional animals were used for baseline assessments. On histology, changes were evident at 32 hours post-treatment. However, RNA-seq analyses revealed that the molecular impact of CPA occurred early within the first 16 hours of exposure to the drug, and that protective effects of AO in combined CPA therapy (CPA+AO) could be achieved during that period. This study demonstrates side effects of CPA on testicular tissue in the first 48 hours on transcriptional scale. Our findings indicate that early administration of AO can mitigate CPA-induced testicular damage in-vivo and suggest that protective treatments against prepubertal testis damage could be developed. However, such protective treatment should not interfere with the desired cytotoxic effects on the malignancies, thus further research is needed to evaluate the timing and how safe administration of the protectant could be achieved for medical treatment.

## INTRODUCTION

Several chemotherapeutic drugs used for treatment of cancer lead to testicular damage and loss of the spermatogonial stem cells (SSCs) (1). SSCs are the precursors of spermatogenesis, a complex and tightly regulated process that involves the proliferation and differentiation of SSCs by which sperm cells are produced in the testis. These undifferentiated cells have the ability to self-renew and differentiate into the various stages of germ cells, give rise to differentiating spermatogonia, which then undergo meiosis to become spermatocytes, spermatids, and ultimately, spermatozoa (2). Cyclophosphamide (CPA) belongs to the alkylating drugs commonly used in cancer treatment protocols that have a demonstrated toxicity effect on the gonads with irreparable damage on SSCs including infertility in cancer survivors.

There are currently no clinical methods available for recovering fertility in adulthood for the individuals treated for cancer in childhood who have developed SSCs loss following a gonadotoxic treatment (3, 4). Since 2003, our team at Karolinska University Hospital has been actively developing and optimizing cryopreservation methods for fertility preservation in children, leading to advancements in protocols that improve the viability and clinical applicability of preserved tissue (5–7). The long-term follow-up of the pre-pubertal boys undergoing gonadal tissue cryopreservation in this prospective cohort has indicated feasibility and low risk of the procedures (6). While cryopreservation of prepubertal ovarian tissues have shown proof of concept, with resumption of hormonal function to induce puberty (8, 9), as well as live births after pregnancy (10, 11), the use of testis tissue cryopreserved has not yet reported success, and there is an urgent need for development of viable fertility preservation options for this patient group (12, 13). Additional shortcomings in male fertility preservation include the lack of clinical biomarkers that could be useful during patient follow-up, and to assess post-treatment effects and severity of testicular damage (6, 12). Given that most pediatric cancer patients recover from the disease, maintaining and preserving fertility will have a major impact on improving their quality of life.

Since there are no comprehensive studies addressing the early toxicity of CPA on testicular tissue (TT) and testicular cells (TC), we aimed to focus on identification of genes and pathways involved in this process. Our goal was to determine the critical and earliest time frame at which CPA impacts TT with SSCs damage to understand the extent of these effects at the gene transcriptional level. Furthermore, as a few publications have suggested that antioxidants (AO) can protect the SSCs, during or under chemotherapy (14), we sought to test if the damage caused by CPA could be reduced by AO when administered concomitantly. Spermatogenesis seems to be sensitive to free radical formation during cancer chemotherapy even in post-pubertal males (15). Previous experimental studies indicate that treatment with the amino acid derivative L-Carnitine (LC) may decrease germ cell apoptosis, and maintain testosterone levels, sperm motility and viability *in-vivo* (16, 17), while N-acetylcysteine (NAC) treatment may reduce DNA damage induced by CPA benefiting sperm histone protamine replacement (15, 18). Additionally, indirect effects to the germ cells mediated through damage of the somatic supportive cell populations may also lead to infertility (1). It would be plausible that a combination of AO could provide a more sustained effect than that obtained with an individual drug (19, 20).

## RESULTS

### Induced damage of CPA treatment on testicular cells and seminiferous tubule morphology is evident in histological assessments

To investigate the morphological changes induced and effects of CPA on prepubertal TT and TCs, we selected 63 pre-pubertal 7-9 postnatal days CBA/B6 F1 mice pups with 3 biological replicates across four treatment arms: 0-hour BT, Ctrl, CPA, AO, and CPA+AO treatment arms. The CPA treatment time marked as start time point of experiment, 0 hour, AO treatments were administered starting at one day before CPA administration and repeated in 24-hour intervals for 3 days (Figure 1A).

**Fig. 1.**
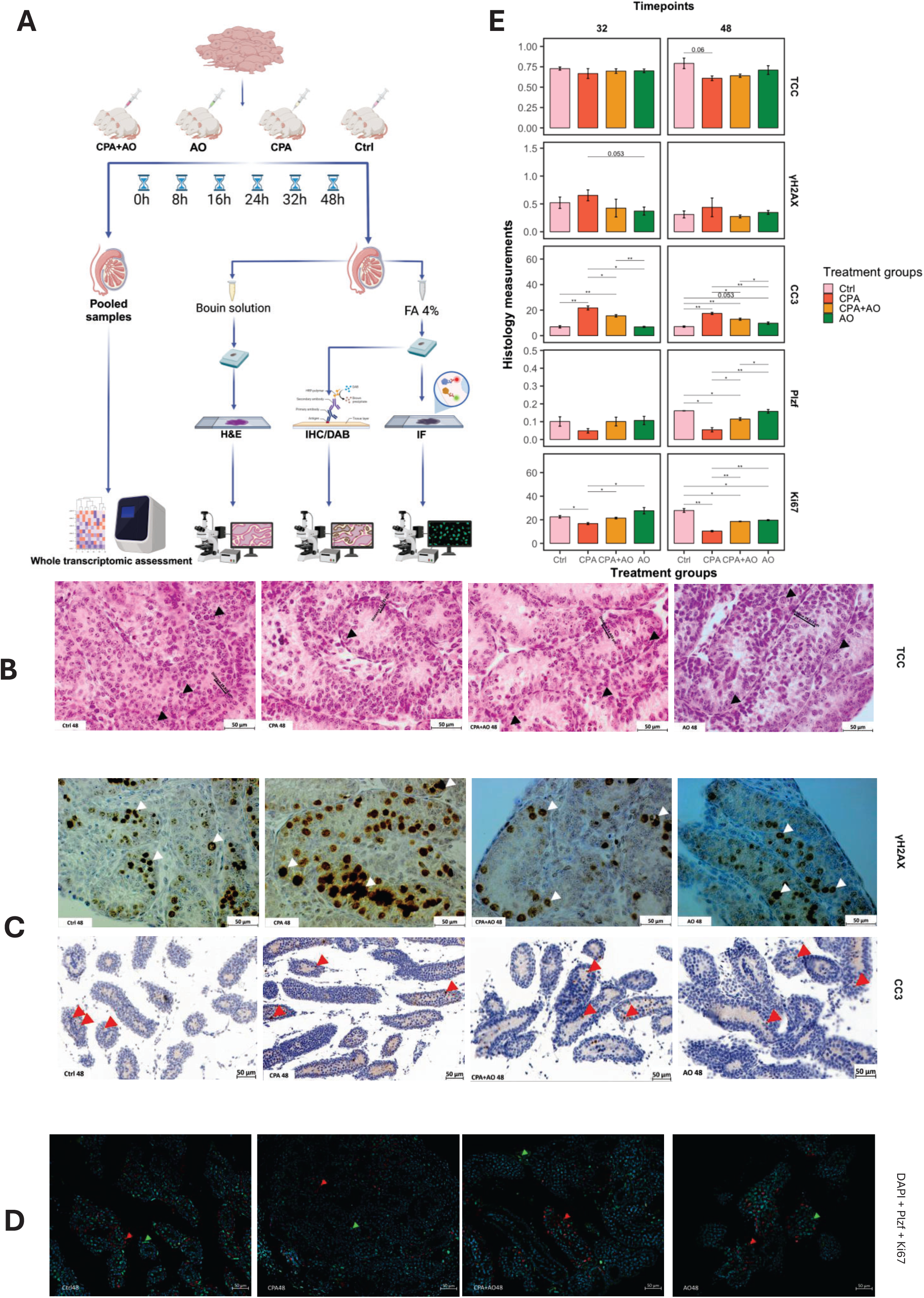
Histological effects of cyclophosphamide with a antioxidant combination on testicular tissue niche in prepubertal mouse models. **A)** Schematic detailing sample preparation and analysis methods used in the study. **B)**, H&E staining testicular tissue at 48 hours after treatment shows the differences between group Ctrl, CPA, CPA+AO and AO in Total cell number per seminiferous tube and cell layers inside the seminiferous tube. **C)**, IHC/DAB staining for γH2AX and CC3 markers for showing the DNA damage and initiation of apoptosis cycle in the groups at 48 hours after CPA treatment. **D)** IF staining for nucleus staining with Dapi (blue); Plzf (green); and Ki67 (red) markers for showing the germ cell numbers inside the seminiferous tube and proliferation activity in the groups at 48 hours after CPA treatment post-sample collection. **E)** Bar plots showing significant differences in histological results for Total cell count/seminiferous area (TCC), γH2AX as DNA damage/repair mechanism ration (DMR) and CC3 positive cells evaluate as a marker of initiation of apoptotic cells and counted as Initiation of apoptosis ration (IAR), *Plzf* positive cells considered as germ cell ration (GCR), while Ki67 positive cells represents cells with high proliferation rate and counted as testicular proliferative capacity ration (TPR) for each treatment group across 32- and 48-hours time points. Ctrl: Control group; CPA: Cyclophosphamide treated group; CPA+AO: Cyclophosphamide combined with antioxidant treatment; AO: Antioxidant treated group; TCC: Total Cell Count in seminiferous tubule; CC3: Cleaved Caspase 3; Plzf: promyelocytic leukaemia zinc finger

Significant loss of germ cells with a thinner layer of cells inside the seminiferous tubes were observed in 48 hours for CPA treated groups (Figure 1B and 1E). A comparison of γH2AX and CC3 shows the difference of DNA damage and apoptosis in CPA treated group in compared to Ctrl and CPA treated groups (Figure 1C and 1E). Plzf positive cells showed no significant change until 32 hours, while the change is evident after 48 hours, similar to Ki67 positive cells reduction at this timepoint (Figure 1D-E). These findings confirm the histological observations and highlight the onset of testicular tissue changes evident on microscopy from 32 hours of CPA administration. We observed morphological changes starting at the 32 hour time point, including the cell lose, reducing the seminiferous tubule cell layers, increasing the apoptosis and DNA damage as well as decrease in proliferation, supporting previous studies of chemotherapy’s effects on TCs and TT (12). Our findings in CPA+AO treated group compared to CPA group suggesting that AO treatment exerts a protective effect on higher TC counts, seminiferous tubule shape and thicker cell layers (Figure 1B-E).

### Effects of antioxidant treatment are evident on testicular cells and testicular tissue in histological assessments

We observed a significant increase of γH2AX formation in the 8 hours and 16 hours samples in the AO treated groups (Supplemental figure 1A-B and supplemental figure 2), without any concomitant reduction the SSCs marker Plzf (Supplemental figure 1C and supplemental figure 2). The AO treatment appears to have induced DNA double-strand breaks, as evidenced by increased γH2AX expression. This DNA damage was accompanied by higher Ki67 levels, suggesting the DNA damage was associated with cellular proliferation. While we observed some degree of apoptosis in the AO-treated testicular samples compared to the control group, these effects were less pronounced than in the CPA-treated group. Importantly, the AO treatment did not appear to impair the stemness of the germ cells, which is expected to recover in subsequent time points.

### The impact of CPA and antioxidant treatments on gene transcription was confirmed following the intervention

Although previous research has documented CPA’s impact on TT and TCs (2, 21, 22), comprehensive studies at the transcriptional level for early effects of CPA remain limited. We performed deep whole-transcriptome RNA-sequencing (RNA-seq) to explore the transcriptional consequences of CPA + AO treatments on TCs, building on the histological observations.

Differential expression (DE) analysis, revealed 42 significantly upregulated genes related to cell death and apoptosis, including *Casp8* and *Erbb2*, in CPA group at 48 hours post-administration. While, 37 genes involved in cell development and spermatogenesis, such *Aifm3, Amhr2, Bik, Ddit3, Gadd45a, Kit, Pak1, Sohlh1* and *Zbtb16* (*Plzf*) were downregulated (Figure 2A). These genes are involved in pathways associated with cellular oxidant detoxification, gonad development, sex differentiation, reproductive system development and reproductive structure development were downregulated compared to the Ctrl group (Figure 2C).

**Fig. 2.**
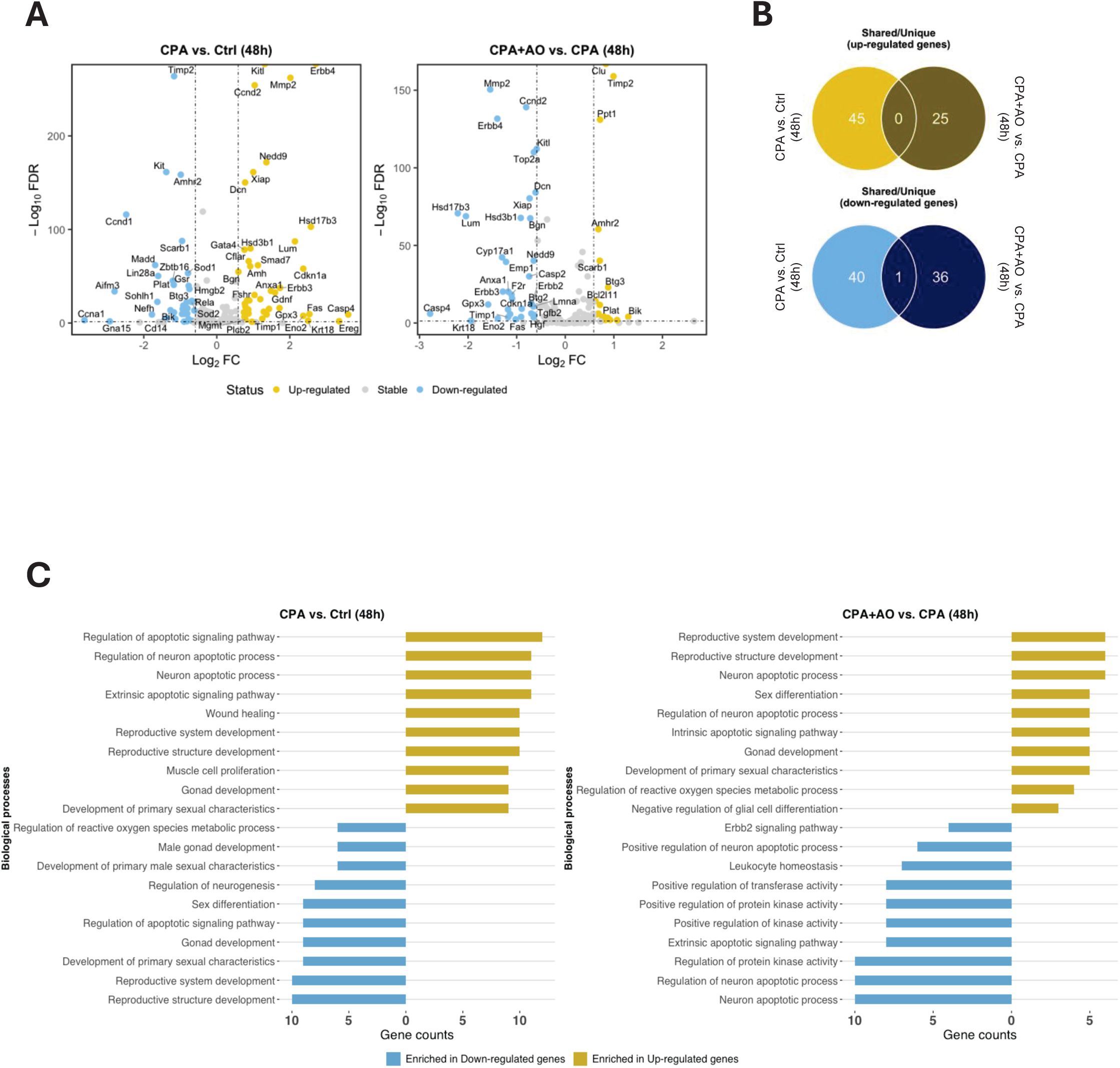
Transcriptional findings of cyclophosphamide effect with antioxidant combination on testicular tissue niche in prepubertal mouse models. Volcano plots showing upregulated (yellow) and downregulated (blue) genes: **A)** CPA group vs. Ctrl at 48 hours, and CPA+AO group vs. CPA group at 48 hours. Genes with fold-change (FC) ≥ 2 and false discovery rate (FDR) < 0.05 are considered significant. **B)** Venn diagrams showing the overlap and uniqueness of upregulated (yellow) and downregulated (blue) genes between comparisons, with the number of differentially expressed genes indicated within each circle. Bar plots of the top 10 enriched biological processes for upregulated and downregulated genes: **C)** CPA vs. Ctrl, and CPA+AO group therapy vs. CPA group at 48 hours. The bar length represents the number of genes contributing to each biological process category. Ctrl: Control group; CPA: Cyclophosphamide treated group; CPA+AO: Cyclophosphamide combined with antioxidant treatment; AO: Antioxidant treated group.

To assess how AO affect CPA caused gene expressional alterations, we compared CPA+AO to CPA treated groups, finding 20 upregulated genes related to cell structure development and spermatogenesis, such as *Atf3* and *Ddit3* (23) (Figure 2A-C). Comparing gene expression changes across these conditions, show no shared upregulated genes, though *Casp2-4, Fas, Gpx3* and *Kitl* genes among 35 downregulated genes, related to extrinsic apoptotic signaling, homeostasis of number of cells and regulation for apoptotic process were the common downregulated genes (Figure 2A-C), additionally, our data shows that there is one shared gene, *CD14*, that is downregulated in this comparison between CPA vs Ctrl to CPA+AO vs CPA groups at 48 hours (Figure 2B). This suggests that the AO in combination with CPA (CPA+AO) treatment could mitigate the detrimental effects of CPA on TCs including apoptosis, decreasing proliferation and total cell and germ cell numbers. A complete list of DE genes and GO analyses of both comparisons are provided in supplemental tables 2 and 3, respectively.

### Early onset of CPA-induced apoptosis is detectable at the transcriptional level

In line with previous studies, we observed the impact of treatments on morphology and histological features, particularly in the induction of cell apoptosis, after 32 hours (24). However, a significant knowledge gap persists regarding the early-stage effects of chemotherapy. To address this, we examined gene expression changes prior to 32 hours, with a particular emphasis on genes linked to apoptosis and cell death, in order to elucidate the early molecular responses to treatment. A comparison of apoptosis gene expression profiles among the treatment groups at different time points revealed distinct gene expression patterns between the Ctrl and CPA groups as early as 16-hour post-treatment (Figure 3A). Principal component analysis (PCA) showed that samples could cluster based on the type of treatment given to the models, except for the CPA group at 16 hours time point (Figure 3B). Notably, a drastic reduction of apoptosis signature activity was observed at 16 hours in CPA treated group while this wasn’t evident in Ctrl group (Figure 3C).

**Fig. 3.**
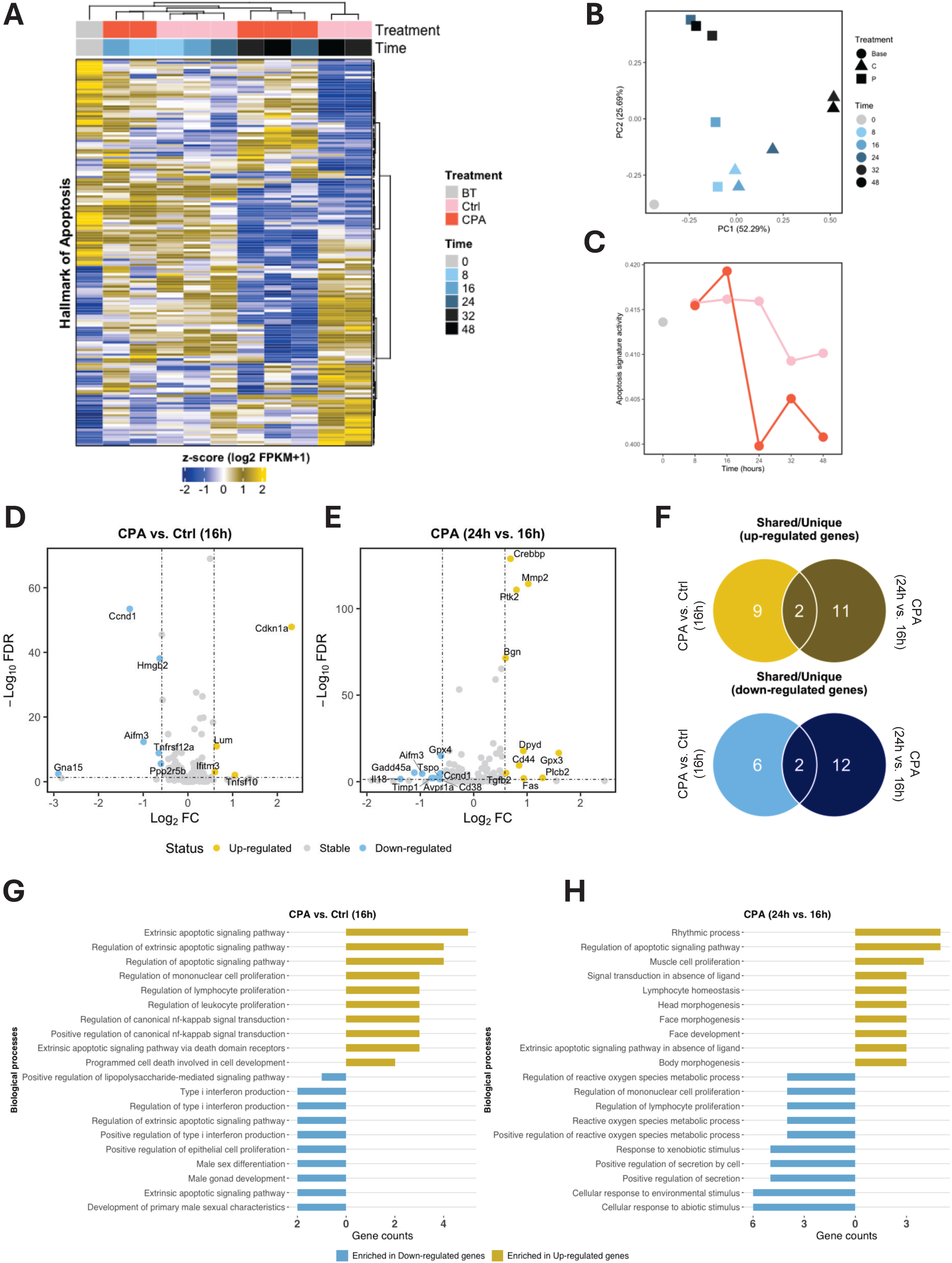
Early effects of cyclophosphamide on apoptosis observed at the transcriptional level. **A)** Heatmap representing the expression profiles of apoptosis-associated genes in Ctrl group and CPA group. Time points are represented by different shades of blue. The control (Ctrl) sample at the base time point (0 hours) is marked as “BT” and shown in grey. **B)** Principal component analysis showing that samples from different treatment groups cluster together, except for the CPA group at 16 hours. **C)** Line plots indicating apoptosis activity in each group at different time points. The BT, Ctrl, and CPA groups are coloured grey, pink, and yellow, respectively. Volcano plots showing upregulated (yellow) and downregulated (blue) genes: **D)** CPA group vs. Ctrl group at 16 hours, and **E)** CPA group at 24 hours vs. 16 hours. Genes with fold-change (FC) ≥ 2 and false discovery rate (FDR) < 0.05 are considered significant. **F)** Venn diagrams showing the overlap and uniqueness of upregulated (yellow) and downregulated (blue) genes between comparisons, with the number of differentially expressed genes indicated in each circle. Bar plots showing the top 10 enriched biological processes for upregulated and downregulated genes: **G)** CPA vs. Ctrl groups at 16 hours, and **H)** CPA group mice at 24 hours vs. 16 hours. The bar lengths represent the number of genes contributing to each biological process category. Ctrl: Control group; CPA: Cyclophosphamide treated group; CPA+AO: Cyclophosphamide combined with antioxidant treatment; AO: Antioxidant treated group.

These findings indicate that significant transcriptional changes emerge as early as 16 hours after treatment. Given the importance of this timepoint, we analyzed the differences in gene expression levels between Ctrl and CPA groups at 16 hours post-treatment to capture the early molecular responses to chemotherapy. Comparing the CPA16 and Ctrl16, we observed up-regulation of genes involved in the extrinsic apoptotic signaling pathway, programmed cell death in cell development, and cytokine-mediated signaling, such as *Cdkn1a*, *Ifitm3* and *Tnfsf10*.

Conversely, genes related to the development of primary male sexual characteristics, male gonad development, male sex differentiation, and type I interferon production regulation, including *Ccnd1*, *GNA15*, *Aifm3* and *Tnfrsf12a*, were downregulated in the CPA16 group compared to the Ctrl16 group (Figure 3D and 2G). These results suggest that the onset of apoptosis, activation of cell death pathways, and disruptions in male gonad development occur earlier than the observed histological changes (after 32 hours of injection in Histology). Therefore, 16 hours time point may represent a critical window for gene expression alterations, providing valuable insights for developing strategies to mitigate the adverse effects of CPA on TCs.

As the drastic drop on apoptosis activity was observed at 24 hours timepoint (Figure 3C), we sought to identify genes that have been impacted at this time. Performing DE analyses between the CPA24 and CPA16 groups revealed up-regulation of *Fas*, *Gpx3*, *Plcb2*, *Ptk2* and *Tgfb2* genes involved in the apoptotic pathway, extrinsic apoptosis signaling pathway, and lymphocyte homeostasis. Conversely, *Aifm3*, *Ccnd*, *Gadd45a*, *il18* and *Timp1* genes that related to reactive oxygen species (ROS), lymphocyte proliferation, and leukocyte proliferation were downregulated in the CPA24 group compared to the CPA16 group (Figure 3E and 3H). These findings indicate that the gene expression changes observed at 16 hours continue to escalate over time, culminating in an exacerbation of apoptosis and cell death signals. These changes may occur through the differential expression of various genes over time, as shown in the Venn diagrams, which indicate that most of the up-or down-regulated genes are unique to the specific time points of 16 or 24 hours (Figure 3F). Our analysis identified *Ereg* and *Plcb2* as genes that were shared upregulated genes, and *Aifm2* and *Ccd1* as genes that were shared downregulated genes in this comparison among CPA vs Ctrl at 16 hours as compared to CPA at 24 hours vs CPA at 16 hours, corroborating the results from the PCA analyses. A complete list of DE genes and GO analyses of both comparisons are provided in supplemental tables 4 and 5, respectively.

### The impact of CPA on testicular cell types and cell proliferation is detectable earlier timepoints at transcriptional level

As we shown in our histological findings (Figure 1 B-D), CPA treatment has detrimental effect on TCs proliferation and numbers after 32-hour of treatment. To examine the early effects of CPA on TCs populations and proliferation prior to 32-hour time point, we conducted transcriptional-level analysis across all treatment groups. Considering our previous findings on CPA’s impact on TT, we anticipated observing the onset of detrimental effects on gene expression by 16 hours post-treatment (2).

The heatmap analysis revealed the expression profiles of the cell proliferation marker Ki67 and key TC types, including Leydig cells (LC), Sertoli cells (SC), and Spermatogonia Stem cells (SSC) in Ctrl and CPA groups. Notably, the gene expression pattern of TC markers in CPA group at 16 hours was distinct from the other samples in this group, supporting that the 16-hour timepoint may also be a critical juncture for transcriptional regulation among the treatment conditions (Figure 4A). PCA analyses based on genes associated with different cell types revealed that samples from the different treatment groups generally clustered together, with the notable exception of the CPA group at the 16-hour timepoint. This suggests that the gene expression profiles of the treatment groups began to diverge starting at the 16-hour timepoint, although prior to that the 8-hour CPA group exhibited a gene expression pattern more similar to the Ctrl group (Figure 4B).

**Fig. 4.**
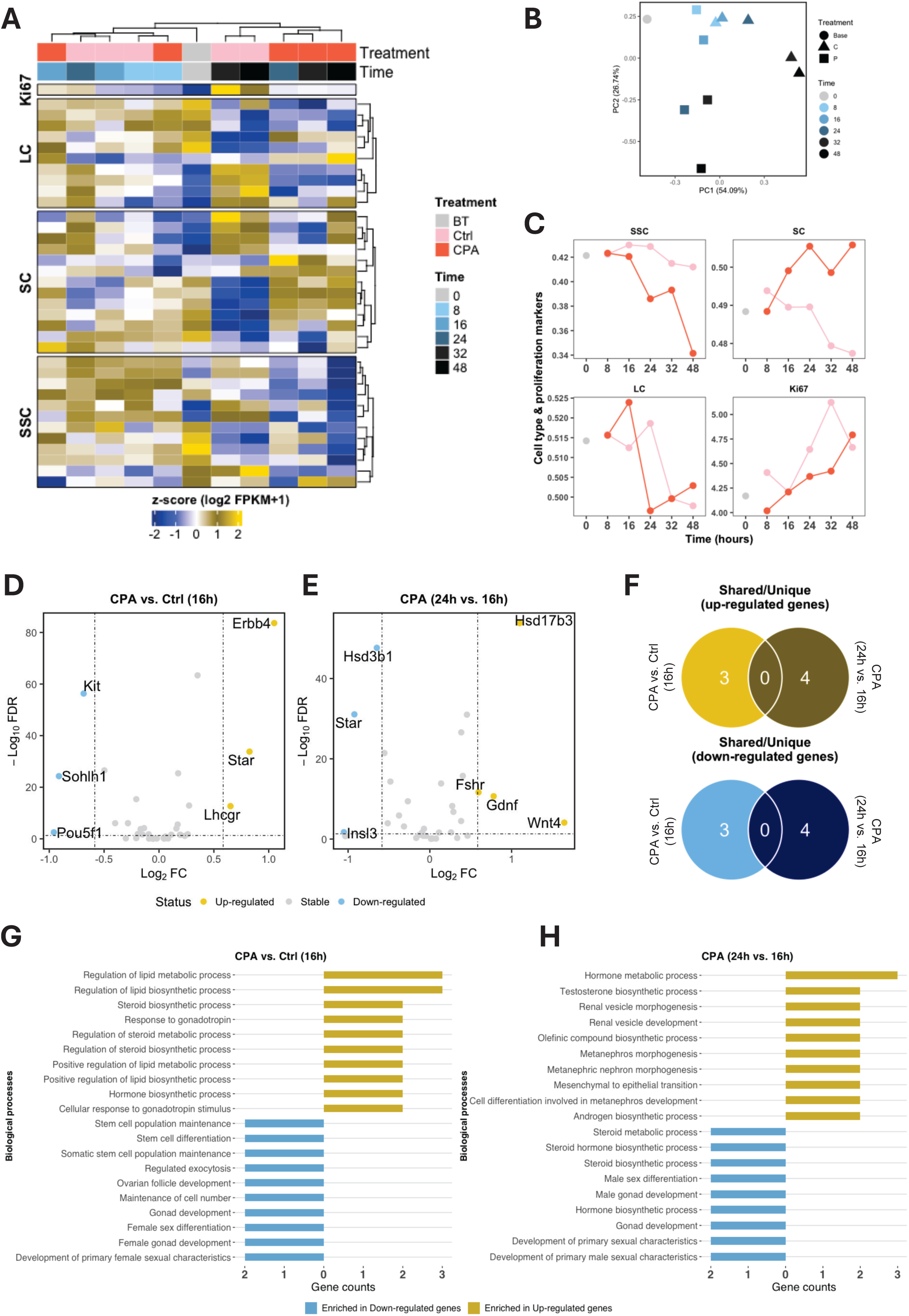
Early effects of cyclophosphamide on cell proliferation and key testicular cells involved in spermatogenesis at the transcriptional level. **A)** Heatmap showing the expression profiles of cell proliferation marker Ki67 and key testicular cells (Leydig cells [LC], Sertoli cells [SC], and Spermatogonial Stem Cells [SSC]) in control and Cyclophosphamide-treated mice. Time points are indicated by different shades of blue, with the control sample at 0 hours marked as “BT” in grey. **B)** Principal Component Analysis illustrating clustering of samples from different treatment groups, except for the CPA group at 16 hours. **C)** Line plots depicting gene expression scores for LC, SC, SSC, and Ki67 across different time points, with colours representing the BT (grey), Ctrl group (pink), and Cyclophosphamide-treated (red) CPA groups. Volcano plot showing upregulated (yellow) and downregulated (blue) genes in **D)** the CPA group compared to Ctrl at 16 hours. And **E)** CPA-treated mice at 24 hours versus 16 hours, highlighting genes with fold-change (FC) ≥ 2 and false discovery rate (FDR) < 0.05. **F)** Venn diagrams displaying overlap and uniqueness of upregulated (yellow) and downregulated (blue) genes between comparisons, with numbers of differentially expressed genes indicated. Bar plots showing the top 10 enriched biological processes for upregulated and downregulated genes **G)** in CPA group versus Ctrl group at 16 hours and **H)** in CPA group at 24 hours compared to 16 hours. Bar lengths represent the number of genes associated with each biological process category. Ctrl: Control group; CPA: Cyclophosphamide treated group; CPA+AO: Cyclophosphamide combined with antioxidant treatment; AO: Antioxidant treated group.

### Testicular cell types exhibit varying recovery capacities after CPA treatment

Further investigation on gene expression patterns of different TCs types, and the proliferation marker Ki67, support the hypothesis that the 16-hour time point is a critical juncture in the response to CPA treatment (Figure 4C). For SSC, the data indicates a significant deviation in gene regulation starting at 16 hours, with no apparent recovery by the end of the experiment.

A similar pattern is observed for SCs, though the divergence between control and treated groups appears to initiate earlier, at 8 hours post-treatment. In contrast, the LC gene expression profile shows differences compared to the other cell types, suggesting this cell population may have a greater capacity for recovery or resistance to the effects of CPA. The pattern of Ki67 expression level changes, illustrates the expected transient impact of a single dose of chemotherapy, with the difference between control and treated groups diminishing by 48 hours (Figure 4C).

### CPA disrupts metabolic pathways and impairs testicular development in the early stages of treatment, before histological changes become apparent

Our investigations on gene expression differences between the CPA and Ctrl groups at 16 hours after treatment reveal upregulation of 3 genes *Erbb4*, *Lhcgr*, and *Star*, which are associated with the regulation of lipid metabolic processes, steroid biosynthetic processes pathways.

Conversely, 3 genes related to the development of primary sexual characteristics, gonad development, and somatic cell population maintenance, including *Kit*, *Pou5f1*, and *Sohlh1*, were downregulated in the CPA compared to the Ctrl group (Figure 4D and G).

Observing the substantial differences in gene expression patterns between the 16 hours and 24 hours CPA treated groups, we further investigated the gene expression changes between these two timepoints. This analysis revealed the upregulation of genes such as *Fshr*, *Gdnf*, *Hsd17b3*, and *Wnt4*, which are involved in testosterone biosynthesis, cellular differentiation, and mesenchymal-to-epithelial transition. Conversely, genes including *Hsd3b1*, *Insl3*, and *Star* associated with gonads development, hormone metabolic processes, and male sexual development and differentiation pathways, were downregulated (Figure 4E and H). However, we did not identify any shared up-or down-regulated genes when comparing CPA to Ctrl at 16 hours, or when comparing CPA at 24 hours to CPA at 16 hours (Figure 4F). A complete list of DE genes and GO analyses of both comparisons are provided in supplemental tables 6 and 7, respectively.

### The early protective effects of antioxidants in combination therapy with CPA are evident at the transcriptional level

Building on our observation that the early effects of chemotherapy are detectable at the gene transcriptional level starting at 16-hour timepoint, whereas morphological changes emerge only after 32 hours, we sought to investigate the protective effects of the chemo-protectant AO, both alone and in combination with CPA therapy (CPA+AO), across various time points.

Profiling the expression of genes associated with TCs types and apoptosis revealed that samples treated with AO and CPA+AO clustered closely with the Ctrl group, supporting the protective effect of AO treatment, whether administered alone or in combination. This indicates that AO treatment could effectively mitigate the impact of CPA on TCs and proliferation pathways (Figure 5A). Furthermore, this pattern is consistent with the analysis of gene signatures related to cell types and apoptotic activity. The AO and CPA+AO groups exhibited lower levels of apoptotic activity and maintained higher expression levels of cell type gene signatures compared to the CPA group, beginning to emerge after the 16-hour time point (Figure 5B).

**Fig. 5.**
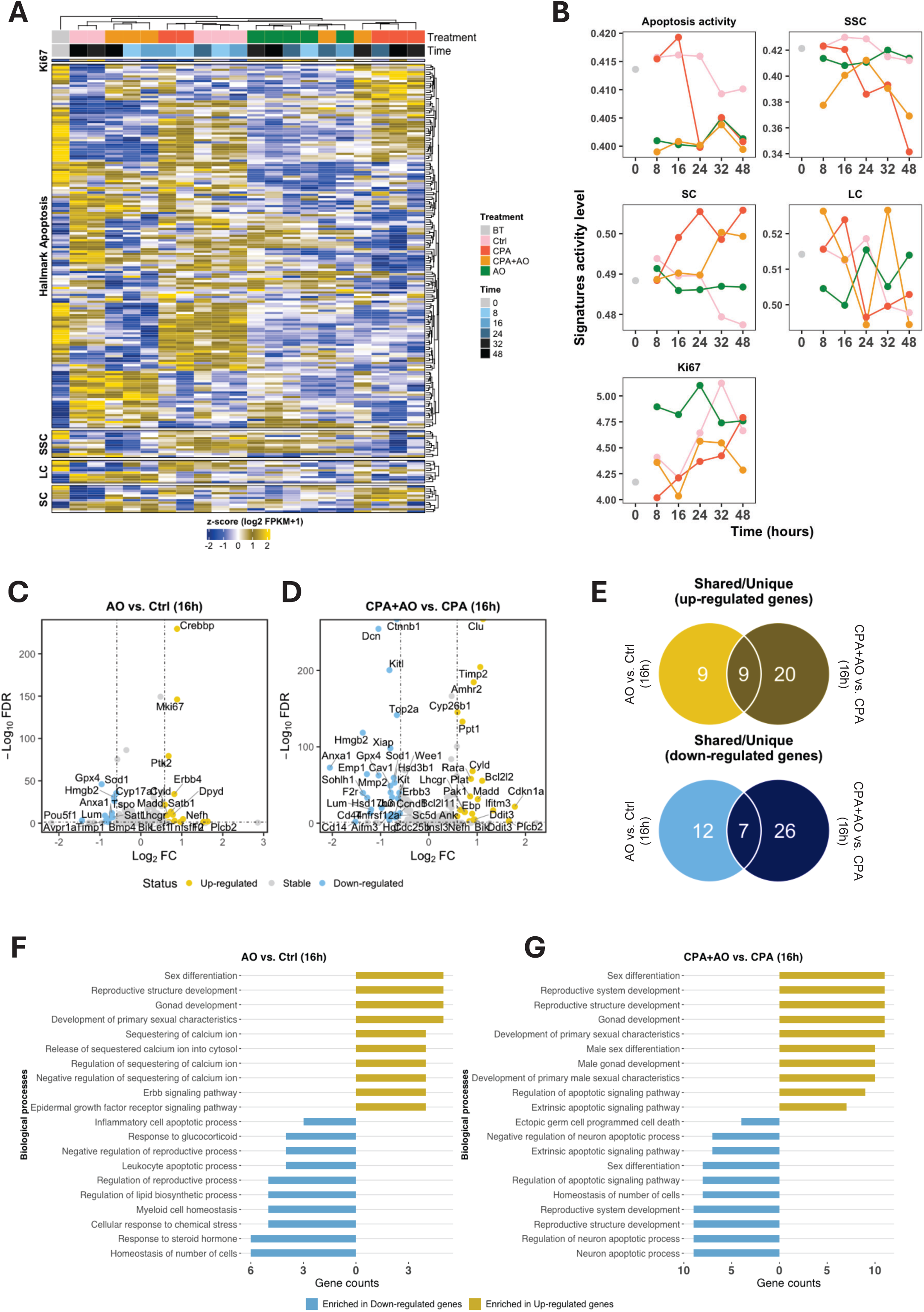
Early benefits of antioxidant treatment on apoptosis, cell proliferation, and key testicular cells involved in spermatogenesis are observed at the transcriptional level. **A)** Heatmap showing expression profiles of apoptosis-associated genes, cell proliferation marker Ki67, and key testicular cells (Leydig cells [LC], Sertoli cells [SC], and Spermatogonial Stem Cells [SSC]) in BT (grey), Ctrl (pink), CPA-treated (red), AO-treated (green), and CPA+AO therapy (yellow) groups. Time points are represented by different shades of blue, with the control sample at 0 hours marked as “BT” in grey. **B)** Line plots illustrating gene expression scores for apoptosis activity, LC, SC, SSC, and Ki67 across different time points, with colours representing the BT (grey), Ctrl (pink), CPA (red), CPA+AO (yellow), and AO (green) groups. Volcano plots showing upregulated (yellow) and downregulated (blue) genes in **C)** the AO group compared to Ctrl at 16 hours and in **D)** CPA+AO group versus CPA group at 16 hours, highlighting genes with fold-change (FC) ≥ 2 and false discovery rate (FDR) < 0.05. **E)** Venn diagrams showing the overlap and uniqueness of upregulated (yellow) and downregulated (blue) genes between comparisons, with the number of differentially expressed genes indicated. Bar plots displaying the top 10 enriched biological processes for upregulated and downregulated genes in **F)** the AO group compared to Ctrl group at 16 hours and in **E)** CPA+AO group versus CPA group at 16 hours. Bar lengths represent the number of genes associated with each biological process category. Ctrl: Control group; CPA: Cyclophosphamide treated group; CPA+AO: Cyclophosphamide combined with antioxidant treatment; AO: Antioxidant treated group.

### Administration of antioxidants positively impacts reproductive development pathways along with CPA treatment

Given the observed differences in apoptosis, cell proliferation, and cell type signature activities between the AO and CPA+AO groups at the 16-hour time point, we conducted DE analysis for AO and CPA+AO groups, along with Ctrl and CPA groups, following 16 hours of treatment. For the AO16 vs. Ctrl16 comparison, 15 genes were found to be upregulated, including *Bik*, *Erbb4*, *Lhcgr*, *Tnfsf10*, and *Madd*, which are involved in gonad development, the epidermal growth factor signaling pathway, and the development of primary sexual characteristics. Conversely, 12 genes, such as *Anxa1*, *Gpx4*, *Pou5f1*, *Tspo*, and *Sat1*, were downregulated, and these are associated with inflammatory cell apoptosis, leukocyte apoptotic process, negative regulation of reproductive processes, and lipid biosynthesis (Figure 5C and F). The DE analyses comparing CPA+AO 16 and CPA16 revealed 23 up-regulated genes, including *Amhr2*, *Bcl2l11*, *Bcl2l2*, *Bik*, *Ddit3*, *Ifitm3*, *Lhcgr*, *Pak1*, and *Timp2*, which are involved in sex differentiation, reproductive structure development, gonad development, and male sex differentiation. Additionally, 29 genes were downregulated, such as *Aifm3*, *Anxa1*, *Ccnd1*, *Kit*, *Sohlh1*, *Tnfrsf12a*, and *Xiap*, that are associated with ectopic germ cell death, gonad development, regulation of apoptotic processes, and homeostasis of cell number (Figure 5D and G). To identify the up-or down-regulated genes in the CPA+AO group that are specifically associated with the protective effects of AOs, even in the presence of chemo toxins, we compared the DE genes between the AO and CPA+AO groups. Our analysis revealed 9 up-regulated genes; *Bik, Cyld, F2, Fasl, Lhcgr, Madd, Nefh, Plcb2 and Tnfsf* and 7 down-regulated genes: *Anxa1, Casp4, Cd14, Gpx4, Hmgb2, Lum and Sod1* shared between the AO and CPA+AO groups, indicating their direct association with the protective effects of AOs despite the presence of CPA in the CPA group (Figure 5E). A complete list of DE genes and GO analyses of both comparisons are provided in supplemental tables 8 and 9, respectively.

## DISCUSSION

This study provides a comprehensive investigation into the early effects of CPA in prepubertal mice using a range of complementary techniques. Notably, our findings demonstrate that CPA, a widely used chemotherapeutic agent, can induce significant transcriptomic changes in the TC as early as 16 hours after injection, and long before any observable histological changes or cellular alterations.

Our histological analysis reveals that CPA treatment induces significant damage on prepubertal TT, which becomes evident approximately 32 hours after administration with a significant impact at 48 hours after treatment. Consistent with previous findings, these effects include a reduction in total cell count, a decline in proliferative activity (25, 26), thinning of cell layers within the seminiferous tubules, degeneration of the seminiferous tubule structure, a decrease in germ cell numbers (27), all of which are characteristic histological findings associated with CPA-induced testicular toxicity following chemotherapy exposure and infertility (2, 28–31). In parallel, we observe a significant increase in apoptosis (CC3) and DNA damage activity (γH2AX). The histological and immunohistochemical analyses revealed that the initial signs of testicular damage, such as reduced germ cell numbers and reduction of proliferation activity, became evident around 32-48 hours after CPA treatment. These findings are consistent with previous studies that have reported similar time frames for the onset of testicular damage following chemotherapy in animal models (32).

Our earlier histological observations at 8 hours and 16 hours time points revealed a significant increase of the DNA damage marker γH2AX in the AO-treated groups, without any concurrent reduction in the proliferation marker Ki67 or the SSCs marker Plzf. This occurred specifically at the 32-hour timepoint after AO administration (8 hours after CPA treatment), supporting our earlier morphological findings on CPA treatment effect. Importantly, this AO-induced effect does not appear to have any prolonged detrimental impacts on TT or TCs. On the contrary, the AO treatment may even confer a protective effect against the side effects of CPA and enhance the recovery rate of both TT and TCs. Notably, our findings suggest that the AO treatment could be a promising approach to mitigate the adverse effects of cancer therapies on male fertility, potentially preserving SSCs function and enabling better reproductive outcomes for prepubertal male cancer patients (14, 15, 17).

Importantly, the transcriptomic analysis provided a more comprehensive understanding of the early molecular events triggered by CPA treatment. Our data showed significant changes in the expression of genes involved in critical processes such as spermatogenesis, apoptosis, and cell cycle control as early as 16 hours after treatment. These early molecular-level changes likely preceded the observable histological and cellular-level alterations, highlighting the importance of utilizing a multifaceted approach to fully understand the impact of chemotherapy on prepubertal TC (21). The identification of these early transcriptional changes is particularly important, as it suggests that the prepubertal testis may be highly sensitive to the damaging effects of chemotherapeutic agents, even at early stages of exposure (33), and it underscores the critical need for the development of targeted interventions and chemo-protective strategies to mitigate the adverse impacts of cancer treatments on future fertility in young patients (34).

The increased expression of cleaved caspase-3 in the prepubertal testes after CPA injection indicates the activation of the apoptosis pathway, which is a critical mechanism by which chemotherapeutic agents can induce testicular damage (35), similarly to how they deplete the ovarian follicles in prepubertal ovaries (36). Consistent with prior studies, we observed an elevated level of γH2AX following CPA treatment, indicating an enhanced DNA damage response that can trigger cell cycle arrest and apoptosis as an early event in chemotherapy-induced testicular toxicity (21). Our findings provide a mechanistic understanding of the pathways leading to germ cell loss and apoptosis in the prepubertal testis.

For further investigation into early events, we use transcriptomic assessment to show the genes and pathways involved behind these morphological changes. Our data demonstrates that CPA exposure results in significant alterations in the expression of genes related to critical testicular development and metabolic pathways, as early as 16 hours post-treatment, well before any morphological changes are observed. The early upregulation of genes like *Erbb4*, *Lhcgr*, and *Star*, involved in the lipid metabolism and steroid biosynthesis, suggesting that CPA may have a direct impact on Leydig cell function at the early stages of alkylating treatment.

In contrast, the downregulation of genes associated with the maintenance of SSCs and the development of germ cells, such as *Kit, Pou5f1*, and *Sohlh1*, suggests that CPA may have a detrimental impact on these critical aspects of testicular function (24). The protective effects of AO therapy, in combination with CPA, are also evident at the transcriptional level, with distinct gene expression patterns emerging as early as 16 hours post-treatment. The upregulation of genes involved in reproductive development pathways, such as *Amhr2, Bcl2l11, Lhcgr,* and *Pak1*, in the AO-treated groups indicates that these compounds can mitigate the deleterious effects of CPA on testicular function and gametogenesis.

Interestingly, the concurrent administration of AO compound partially alleviated the detrimental effects of CPA on the TC, as evidenced by the upregulation of relevant pathways. We suggest using a combination of LC and NAC as both are FDA-approved and proposed antioxidants for protecting testicular cells against cyclophosphamide treatment. For enhancing the protectant effect of these AOs and decreasing negative side effects of the higher dosage usage. Our findings have important implications for the management and monitoring of prepubertal cancer patients undergoing chemotherapy. The early molecular changes identified in this study may serve as potential biomarkers or targets for the development of interventions aimed at mitigating the detrimental effects of CPA on testicular function and fertility preservation (37). These findings could lead to the development of more targeted and effective strategies to protect the delicate prepubertal TT from the damaging effects of chemotherapeutic agents, ultimately preserving fertility and improving the overall well-being of young cancer patients.

Our study has some limitations. First, as a preclinical investigation, it does not yet extend to human samples, particularly those related to TCs, fertility, and immune cell interactions and their impact on normal function of TCs. Additionally, while we have provided insight into the overall transcriptomic changes induced by CPA using bulk RNA-seq, a more granular approach, such as single-cell analysis, could help to uncover the specific effects on distinct cell populations within the TT. This would provide a clearer understanding of how CPA affects various cell types, including SSCs and immune cells, and help elucidate mechanisms that may contribute to long-term fertility risks and testis tissue damage.

Overall, our findings demonstrate the utility of transcriptional profiling to elucidate the early molecular events underlying CPA-induced testicular damage and the protective effects of AO therapy, which may inform the development of targeted strategies to preserve fertility in pediatric cancer patients (24).

### Conclusions

This study demonstrates that CPA exposure at the prepubertal age can have early and severe consequences on testicular development and function, including damage to the germ cell population and disruption of key metabolic and steroidogenic pathways. The findings highlight the importance of early intervention with antioxidants combination therapies, such as LC and NAC, to help preserve testicular integrity and potentially mitigate the late effects of treatment of cancer in childhood due to gonadal toxicity. The early molecular changes identified in this study may serve as potential biomarkers for the development of interventions aimed at protecting the delicate prepubertal TT from the damaging effects of chemotherapeutic agents.

## MATERIALS AND METHODS

### In-Vivo Mice Models and treatment arms

In this research 7-9 days old CBA/B6 F1 prepubertal mice were used. The mature animals (B6 males and CBA females mated for producing the pups) were obtained from the animal research facility of Karolinska Hospital (Huddinge, Sweden), and kept in animal houses under standard conditions (at 22 ± 2 °C room temperature, 50–55% humidity, light/dark cycles: 12/12 h per day). Sixty pre-pubertal mice were randomly assigned to 1) intraperitoneal treatment with 100 mg/Kg CPA alone; 2) CPA combined with AO (CPA+AO) L-Carnitine (LC 50mg/kg) and N-acetyl cysteine (NAC 50 mg/kg); 3) AO alone; and 4) control receiving saline solution (Ctrl) (4 groups; 5 timepoints; 3 replicas; in total N=15 each group); 3 additional pups were used for base time point analyses control group (BT) (in total: 63). All animal procedures were approved by Karolinska Institute and the regional ethics committee for animal research in accordance with the Animal Protection Law, the Animal Protection Regulation, the Regulation of the Swedish National Board for Laboratory Animals, identified Dnr 1372 (date: January 24, 2018).

Based on our preliminary results for this study and previous research (38, 39) we used a high single dose of CPA (100 mg/kg) to cause testicular damage by intraperitoneal injection (IP). The AO treatment included a combination of L-carnitine (LC 50mg/kg), and N-Acetyl cysteine (NAC 50 mg/kg) solved in DPBS that were initiated 24 hours before the CPA injection and repeated at 24 hours intervals for 3 days.

### Sampling

The testes were collected at the specified timepoints (0, 8, 16, 24, 32, 48 hours after CPA injection) following cervical dislocation and one testis from each animal was bisected. From each testis, one half was fixed in Bouin’s solution for histological analysis via Hematoxylin and Eosin (H&E) staining, while the other half was fixed in 4% paraformaldehyde for immunohistochemical and immunofluorescence studies. The contralateral testes were removed and rapidly frozen for whole transcriptomic analysis (Figure 1A). The first 3 pups sacrificed at the start point CPA treatment as base time point group (BT), then at 8, 16, 24, 32, 40, and 48 hours after CPA administration, 3 mice from each group were euthanized, and their testes were dissected.

### Histology and Immunohistochemical methods

#### 1) H&E staining

The tissue samples were fixed in Bouin’s solution, processed using an automated system, and embedded in paraffin. Five-micrometer thick sections were mounted on glass slides and heat-treated for 1-3 hours at 56°C. After deparaffinization in xylene and rehydration through a graded alcohol series, the sections were stained with Mayer’s hematoxylin for 5 minutes followed by 0.2% Eosin Y for 75 seconds. Rapid dehydration through graded alcohols, xylene treatment, and cover slipping with Pertex medium were then performed.

H&E-stained sections of each sample were evaluated, and micrographs were captured using an optical microscope. Identification of cell types, niche degeneration, and the morphology of seminiferous tubules were assessed. Gonocytes lining the basal membrane were recognized by their large, round nuclei and distinct cell borders, while undifferentiated spermatogonia were identified by their characteristic oval nuclei with few nucleoli. H&E staining was performed for histological assessment and total cell number count ration for seminiferous areas (TCC ratio).

#### 2) IHC/DAB staining

The formalin-fixed samples underwent automated tissue processing and were embedded in paraffin (FFPE). Five-micrometer-thick FFPE sections were mounted on glass slides and heat-treated for 3 hours at 56°C. After deparaffinization in xylene and rehydration in alcohol, a heat-induced epitope retrieval was performed using a Decloaking Chamber set at 110°C for 5 minutes in Citrate buffer at pH 6. To quench endogenous peroxidase activity, a 30-minute incubation in 0.15% hydrogen peroxide was carried out at room temperature, followed by a 15-minute blocking step using BioCare Medical Background Sniper. The primary antibodies were diluted in 1% BSA as follows: CC3 (Thermofisher, PA5-77887, 1:200), Plzf (Abcam, ab189849, 1:200), Ki67 (Novus, NB500-170, 1:100) and γH2AX (Thermofisher, BS-37779R 1:200). Incubation with the primary antibodies occurred overnight at 4°C in a humid chamber. For detection, the BioCare Medical Mach-1 Universal HRP-Polymer Kit was used according to the protocol, with a 5-minute DAB reaction time. The sections were counterstained with Mayer’s hematoxylin for 1 minute, followed by dehydration through graded alcohols, xylene treatment, and cover slipping with Pertex. Immunohistochemical staining (IHC/DAB) was performed and γH2AX positive cells counted as surrogate of DNA damage and counted as DNA damage/repair mechanism ration (DMR), along with Cleaved Caspase 3 (CC3) positive cells evaluate as a marker of initiation of apoptotic cells and counted as initiation of apoptosis ratio (IAR) (Figure 1C and 1E).

#### 3) IF Staining

The initial preparation steps were identical to those used for FFPE samples in IHC/DAB, followed by the use of a Decloaking Chamber set at 110°C for 5 minutes in Citrate buffer at pH 6. Permeabilization was achieved by a 30-minute incubation in 0.2% Triton X-100, followed by a 60-minute blocking step in a solution containing 20% normal goat serum and 5% BSA in PBS. The primary antibodies, Ki67 (Cellsignaling, CST 9129T, 1:400) and Plzf (Santa Cruz sc-28319, 1:200), were diluted in 1% BSA. Incubation with the primary antibodies occurred overnight at 4°C in a humid chamber. For visualization, a mixture of Alexa Fluor anti-rabbit 488 (Thermo Scientific A11034, 1:1000) and Alexa Fluor anti-mouse 555 (Thermo Scientific A32727, dil 1:10000) were applied at room temperature for 30 minutes. The sections were then cover slipped using VECTASHIELD Antifade Mounting Medium with DAPI. To ensure consistent quantification, the number of Ki67-, Plzf-, and DAPI-positive nuclei were counted using the QuPath Cell Analyzer. The proportion of Ki67-positive to Dapi-positive cells was calculated as the ratio between Ki67-positive nuclei and Dapi-positive nuclei on each sample section, representing the proliferation index of each sample. Similarly, the proportion of Plzf-positive to Dapi-positive cells was calculated as the ratio between Plzf-positive nuclei and Dapi-positive nuclei on each sample section, representing the germ cell index of each sample. Furthermore, immunofluorescence staining was performed to identify promyelocytic leukemia zinc finger (Plzf) positive cells considered as germ cell ration (GCR), while Ki67 positive cells represents cells with high proliferation rate and counted as testicular proliferative capacity ration (TPR) (Figure 1D, 1E).

### RNA extraction method

Total RNA was extracted from mouse testis tissue using the EchoLUTION Tissue RNA Kit (50) (BioEcho Life Sciences, Germany) following the manufacturer’s protocol. Briefly, approximately 30 mg of freshly isolated or frozen testis tissue was homogenized using a mechanical homogenizer in 450 µL of Lysis Buffer R provided with the kit. Homogenization was performed on ice to preserve RNA integrity.

Following homogenization, the lysate was incubated at room temperature for 5 minutes to ensure complete cell lysis and nucleoprotein dissociation. To enhance RNA purity and eliminate genomic DNA contamination, the lysate was processed through the EchoLUTION Spin Column system. This single-step purification method employs selective binding of contaminants to the matrix while allowing pure RNA to flow through the column.

The RNA was eluted in 50 µL of RNase-free water. RNA quantification and integrity were assessed with an Agilent 5400 Bioanalyzer (Agilent Technologies, USA), yielding an RNA Integrity Number (RIN) consistently above 8.0, confirming high-quality RNA suitable for downstream applications. The extracted RNA was stored at −80°C for future analysis.

### Whole transcriptome RNA-seq quality control and alignment

RNA-sequencing was performed on pooled biological triplicates from each treatment group, utilizing the NovaSeq X Plus platform with sequencing depth of 50X. Quality control of the raw sequencing data was conducted using FastQC, with reads containing adapter contamination, more than 10% ambiguous nucleotides (N > 10%), or over 50% low-quality bases (Phred score < 5) being removed. For alignment, HISAT2 (40), a graph-based aligner designed for efficient and sensitive mapping of RNA-seq reads, was employed. HISAT2 uses a hierarchical index strategy, leveraging both a global GFM index and local GFM indexes to improve the alignment of reads spanning multiple exons, thereby increasing alignment accuracy and sensitivity. The reads were aligned to the Mus musculus reference genome (GRCm39).

### Gene expression quantification and normalization

Transcript abundance, directly proportional to gene expression, gene length, and sequencing depth, was normalized using Fragments Per Kilobase of transcript per Million mapped reads (FPKM) accounted for both gene length and sequencing depth (41).

### Differential expression and gene ontology analyses

Differential expression analysis was performed using edgeR (v. 4.2.1) in R (42), with the TMM (Trimmed Mean of M-values) normalization method applied. False-positive discovery rate (FDR) was calculated using Benjamini-Hochberg correction. Gene Ontology (GO) enrichment analysis was conducted using the enrichGO function from the ClusterProfiler package (v. 4.12.6) in R (43).

### Gene signature development

Apoptosis signature activity was generated by consolidating the expression levels of genes associated with the hallmark of apoptosis, as retrieved from the Human MSigDB database (44). The gene sets for various cell types, including Leydig cells (LC), Sertoli cells (SC), and Spermatogonial stem cells (SSCs), were retrieved from previous studies (45, 46). A complete list of genes is provided in Supplemental table 1.

### Statistical analysis

For differential expression analysis, genes meeting the criteria of fold change ≥1.5 and FDR ≤ 0.05 were considered significant. GO terms associated with differentially expressed genes were considered significant at FDR < 0.05. For other comparisons FDRs were calculated unless otherwise indicated.

## Funding

The Swedish Childhood Cancer Foundation (PR2016-0115, PR2020-0136)

The Swedish Cancer Society (CAN 2017/704, 20 0170 F)

The Swedish Research Council (Dnr 2020-02230)

Radiumhemmets Research Funds Grant for clinical researchers 2020-2025

The Stockholm County Council (FoUI-953912)

The Karolinska institute Research grants in pediatrics from the Birgitta and Carl-Axel Rydbeck Donation, 2020-00339 to KARW.

The SciLifeLab & Wallenberg Data Driven Life Science Program (KAW 2024.0159) to AL.

## Author contributions

Conceptualization: AE, ARP, KAR-W

Methodology: AE, ARP, AL, KAR-W

Investigation: AE

Sampling: AE, XH, RM

Visualization: AE, AL

Funding acquisition: KAR-W

Project administration: AE

Supervision: AL, KAR-W

Writing – original draft: AE, AL

Writing – review & editing: AE, ARP, AL, KAR-W

## Data and materials availability

All data are available in the main text or the supplementary materials, R scripts to reproduce the main figures of this study are (will be) available at: https://github.com/arianlundberg/EarlyCpEffects.

## List of Supplementary Materials

**Fig. S1. Effect of cyclophosphamide and antioxidant treatment on prepubertal mice testicular tissue and cells.**

**A)**, IHC/DAB staining for γH2AX marker for showing the DNA damage or double stranding due to proliferation of cells among the groups at base time point compared to 8- and 16-hours after CPA treatment. **B)**, IHC/DAB staining for *Plzf* gene marker for showing the germ cells numbers inside the seminiferous tube in different the groups at base time point, 8- and 16-hours after CPA treatment.

**Fig. S2. Effect of cyclophosphamide and antioxidant treatment on prepubertal mice testicular tissue and cells and comparison between groups.**

Bar plots showing the differences in histological results for Total cell count/seminiferous area (TCC), γH2AX, CC3 and Ki67 marker for comparing the DNA damage, apoptosis, germ cell number and proliferation activity changes and comparing the treatment groups at 8-, 16-, and 24 hours after treatment.

**Table S1.** List of genes associated with apoptosis, cell proliferation, and key testicular cells involved in spermatogenesis included in the study.

**Table S2.** List of upregulated and downregulated genes in mice treated with Cyclophosphamide compared to control and in mice treated with combination therapy compared to Cyclophosphamide at 48 hours.

**Table S3.** List of enriched biological processes associated with upregulated and downregulated genes in mice treated with Cyclophosphamide compared to control and in mice treated with combination therapy compared to Cyclophosphamide at 48 hours.

**Table S4.** List of upregulated and downregulated genes in mice treated with antioxidants compared to control at 48 hours.

**Table S5.** List of enriched biological processes associated with upregulated and downregulated genes in mice treated with antioxidants compared to control at 48 hours.

**Table S6.** List of upregulated and downregulated apoptosis-associated genes in control mice and Cyclophosphamide-treated mice.

**Table S7.** List of enriched biological processes associated with upregulated and downregulated apoptosis-associated genes in control mice and Cyclophosphamide-treated mice.

**Table S8.** List of upregulated and downregulated genes associated with the cell proliferation marker Ki67 and key testicular cells (Leydig cells [LC], Sertoli cells [SC], and Spermatogonial Stem Cells [SSC]) in control and Cyclophosphamide-treated mice.

**Table S9.** List of enriched biological processes associated with upregulated and downregulated genes related to the cell proliferation marker Ki67 and key testicular cells (Leydig cells [LC], Sertoli cells [SC], and Spermatogonial Stem Cells [SSC]) in control and Cyclophosphamide-treated mice.

**Table S10.** List of upregulated and downregulated genes associated with apoptosis, the cell proliferation marker Ki67, and key testicular cells (Leydig cells [LC], Sertoli cells [SC], and Spermatogonial Stem Cells [SSC]) in control and antioxidant-treated mice, as well as in mice treated with combination therapy versus Cyclophosphamide at 16 hours.

**Table S11.** List of enriched biological processes associated with upregulated and downregulated genes related to apoptosis, the cell proliferation marker Ki67, and key testicular cells (Leydig cells [LC], Sertoli cells [SC], and Spermatogonial Stem Cells [SSC]) in control and antioxidant-treated mice, as well as in mice treated with combination therapy versus Cyclophosphamide at 16 hours.

## References and Notes

1. Anderson RA, Mitchell RT, Kelsey TW, Spears N, Telfer EE, Wallace WHB. Cancer treatment and gonadal function: experimental and established strategies for fertility preservation in children and young adults. The Lancet Diabetes & Endocrinology. 2015;3(7):556–67.

2. Stukenborg JB, Jahnukainen K, Hutka M, Mitchell RT. Cancer treatment in childhood and testicular function: the importance of the somatic environment. Endocr Connect. 2018;7(2):R69–R87.

3. Cui Y, Harteveld F, Ba Omar HAM, Yang Y, Bjarnason R, Romerius P, et al. Prior exposure to alkylating agents negatively impacts testicular organoid formation in cells obtained from childhood cancer patients. Human reproduction open. 2024;2024(3):hoae049.

4. Duffin K, Neuhaus N, Andersen CY, Barraud-Lange V, Braye A, Eguizabal C, et al. A 20-year overview of fertility preservation in boys: new insights gained through a comprehensive international survey. Human reproduction open. 2024;2024(2):hoae010.

5. Rodriguez-Wallberg KA, Marklund A, Lundberg F, Wikander I, Milenkovic M, Anastacio A, et al. A prospective study of women and girls undergoing fertility preservation due to oncologic and non-oncologic indications in Sweden–Trends in patients’ choices and benefit of the chosen methods after long-term follow up. Acta obstetricia et gynecologica Scandinavica. 2019;98(5):604–15.

6. Borgström B, Fridström M, Gustafsson B, Ljungman P, Rodriguez-Wallberg KA. A prospective study on the long-term outcome of prepubertal and pubertal boys undergoing testicular biopsy for fertility preservation prior to hematologic stem cell transplantation. Pediatric Blood & Cancer. 2020;67(9):e28507.

7. Wikander I, Lundberg FE, Nilsson H, Borgström B, Rodriguez-Wallberg KA. A Prospective Study on Fertility Preservation in Prepubertal and Adolescent Girls Undergoing Hematological Stem Cell Transplantation. Frontiers in Oncology. 2021:2560.

8. Donnez J, Dolmans M-M, Pellicer A, Diaz-Garcia C, Serrano MS, Schmidt KT, et al. Restoration of ovarian activity and pregnancy after transplantation of cryopreserved ovarian tissue: a review of 60 cases of reimplantation. Fertility and sterility. 2013;99(6):1503–13.

9. Missontsa MM, Bernaudin F, Fortin A, Dhédin N, Pondarré C, Yakouben K, et al. Ovarian tissue cryopreservation for fertility preservation before hematopoietic stem cell transplantation in patients with sickle cell disease: safety, ovarian function follow-up, and results of ovarian tissue transplantation. Journal of Assisted Reproduction and Genetics. 2024;41(4):1027–34.

10. Matthews SJ, Picton H, Ernst E, Andersen CY. Successful pregnancy in a woman previously suffering from β-thalassemia following transplantation of ovarian tissue cryopreserved before puberty. Minerva ginecologica. 2018;70(4):432–5.

11. Rodriguez-Wallberg KA, Tanbo T, Tinkanen H, Thurin-Kjellberg A, Nedstrand E, Kitlinski ML, et al. Ovarian tissue cryopreservation and transplantation among alternatives for fertility preservation in the Nordic countries–compilation of 20 years of multicenter experience. Acta obstetricia et gynecologica Scandinavica. 2016;95(9):1015–26.

12. Lopes F, Tholeti P, Adiga SK, Anderson RA, Mitchell RT, Spears N. Chemotherapy induced damage to spermatogonial stem cells in prepubertal mouse in vitro impairs long-term spermatogenesis. Toxicology reports. 2021;8:114–23.

13. Goossens E, Jahnukainen K, Mitchell R, Van Pelt A, Pennings G, Rives N, et al. Fertility preservation in boys: recent developments and new insights. Human reproduction open. 2020;2020(3):hoaa016.

14. Ghafouri-Fard S, Shoorei H, Abak A, Seify M, Mohaqiq M, Keshmir F, et al. Effects of chemotherapeutic agents on male germ cells and possible ameliorating impact of antioxidants. Biomedicine & Pharmacotherapy. 2021;142:112040.

15. Shittu SA, Shittu S-T, Akindele OO, Kunle-Alabi OT, Raji Y. Protective action of N-acetylcysteine on sperm quality in cyclophosphamide-induced testicular toxicity in male Wistar rats. JBRA assisted reproduction. 2019;23(2):83.

16. Zhu B, Zheng Y-f, Zhang Y-y, Cao Y-s, Zhang L, Li X-g, et al. Protective effect of L-carnitine in cyclophosphamide-induced germ cell apoptosis. Journal of Zhejiang University-SCIENCE B. 2015;16(9):780–7.

17. Cao Y, Wang X, Li S, Wang H, Yu L, Wang P. The effects of l-carnitine against cyclophosphamide-induced injuries in mouse testis. Basic & clinical pharmacology & toxicology. 2017;120(2):152–8.

18. Ni K, Spiess AN, Schuppe HC, Steger K. The impact of sperm protamine deficiency and sperm DNA damage on human male fertility: a systematic review and meta-analysis. Andrology. 2016;4(5):789–99.

19. Wei G, Zhou Z, Cui Y, Huang Y, Wan Z, Che X, et al. A Meta-Analysis of the Efficacy of L-Carnitine/L-Acetyl-Carnitine or N-Acetyl-Cysteine in Men With Idiopathic Asthenozoospermia. American Journal of Men’s Health. 2021;15(2):15579883211011371.

20. Owumi SE, Akomolafe AP, Imosemi IO, Odunola OA, Oyelere AK. N-acetyl cysteine co-treatment abates perfluorooctanoic acid-induced reproductive toxicity in male rats. Andrologia. 2021;53(5):e14037.

21. Allen CM, Lopes F, Mitchell RT, Spears N. How does chemotherapy treatment damage the prepubertal testis? Reproduction. 2018;156(6):R209–R33.

22. Allen CM, Lopes F, Mitchell RT, Spears N. Comparative gonadotoxicity of the chemotherapy drugs cisplatin and carboplatin on prepubertal mouse gonads. Molecular human reproduction. 2020;26(3):129–40.

23. Alonso CA, David CD, Toufaily C, Wang Y, Zhou X, Ongaro L, et al. Activating transcription factor 3 stimulates follicle-stimulating hormone-β expression in vitro but is dispensable for follicle-stimulating hormone production in murine gonadotropes in vivo. Endocrinology. 2023;164(5):bqad050.

24. Smart E, Lopes F, Rice S, Nagy B, Anderson RA, Mitchell RT, et al. Chemotherapy drugs cyclophosphamide, cisplatin and doxorubicin induce germ cell loss in an in vitro model of the prepubertal testis. Scientific reports. 2018;8(1):1–15.

25. Medrano JV, Hervás D, Vilanova-Pérez T, Navarro-Gomezlechon A, Goossens E, Pellicer A, et al. Histologic Analysis of Testes from Prepubertal Patients Treated with Chemotherapy Associates Impaired Germ Cell Counts with Cumulative Doses of Cyclophosphamide, Ifosfamide, Cytarabine, and Asparaginase. Reproductive Sciences. 2021;28(2):603–13.

26. Jahnukainen K, Ehmcke J, Hou M, Schlatt S. Testicular function and fertility preservation in male cancer patients. Best practice & research Clinical endocrinology & metabolism. 2011;25(2):287–302.

27. Stukenborg J-B, Alves-Lopes J, Kurek M, Albalushi H, Reda A, Keros V, et al. Spermatogonial quantity in human prepubertal testicular tissue collected for fertility preservation prior to potentially sterilizing therapy. Human Reproduction. 2018;33(9):1677–83.

28. Masliukaite I, Hagen JM, Jahnukainen K, Stukenborg J-B, Repping S, van der Veen F, et al. Establishing reference values for age-related spermatogonial quantity in prepubertal human testes: a systematic review and meta-analysis. Fertility and Sterility. 2016;106(7):1652–7. e2.

29. Portela JM, Heckmann L, Wistuba J, Sansone A, van Pelt AM, Kliesch S, et al. Development and disease-dependent dynamics of spermatogonial subpopulations in human testicular tissues. Journal of Clinical Medicine. 2020;9(1):224.

30. Medrano JV, Hervás D, Vilanova-Pérez T, Navarro-Gomezlechon A, Goossens E, Pellicer A, et al. Histologic Analysis of Testes from Prepubertal Patients Treated with Chemotherapy Associates Impaired Germ Cell Counts with Cumulative Doses of Cyclophosphamide, Ifosfamide, Cytarabine, and Asparaginase. Reprod Sci. 2021;28(2):603–13.

31. Masliukaite I, Ntemou E, Feijen EA, van de Wetering M, Meissner A, Soufan AT, et al. Childhood cancer and hematological disorders negatively affect spermatogonial quantity at diagnosis: a retrospective study of a male fertility preservation cohort. Human Reproduction. 2023:dead004.

32. Anan HH, Wahba NS, Abdallah MA, Mohamed DA. Histological and immunohistochemical study of cyclophosphamide effect on adult rat testis. Int J Sci Rep. 2017;3(2):39–48.

33. Tharmalingam MD, Matilionyte G, Wallace WHB, Stukenborg JB, Jahnukainen K, Oliver E, et al. Cisplatin and carboplatin result in similar gonadotoxicity in immature human testis with implications for fertility preservation in childhood cancer. BMC Med. 2020;18(1):374.

34. Delessard M, Saulnier J, Rives A, Dumont L, Rondanino C, Rives N. Exposure to chemotherapy during childhood or adulthood and consequences on spermatogenesis and male fertility. International journal of molecular sciences. 2020;21(4):1454.

35. Mohamed RH, Karam RA, Hagrass HA, Amer MG, Abd El-Haleem MR. Anti-apoptotic effect of spermatogonial stem cells on doxorubicin-induced testicular toxicity in rats. Gene. 2015;561(1):107–14.

36. Hao X, Reyes Palomares A, Anastácio A, Liu K, Rodriguez-Wallberg KA. Evidence of apoptosis as and early event leading to cyclophosphamide-induced primordial follicle depletion in a prepubertal mouse model. Frontiers in Endocrinology. 2024;15:1322592.

37. Mittra I, Pal K, Pancholi N, Shaikh A, Rane B, Tidke P, et al. Prevention of chemotherapy toxicity by agents that neutralize or degrade cell-free chromatin. Annals of Oncology. 2017;28(9):2119–27.

38. Eskafinoghani A, Hao X, Palomares AR, Rodriguez-Wallberg KA. OPTIMIZING THE TESTIS TISSUE CULTURE POST CYCLOPHOSPHAMIDE TREATMENT BY COMBINED ANTIOXIDANT SUPPLEMENTS Fertility and Sterility. 2023;120(4):e9.

39. Hao X, Anastacio A, Vinals-Ribe L, Santamaria Lacuesta A, Diakaki C, Alonso de Mena S, et al. Follicle Rescue From Prepubertal Ovaries After Recent Treatment With Cyclophosphamide-An Experimental Culture System Using Mice to Achieve Mature Oocytes for Fertility Preservation. Front Oncol. 2021;11:682470.

40. Kim D, Langmead B, Salzberg SL. HISAT: a fast spliced aligner with low memory requirements. Nature methods. 2015;12(4):357–60.

41. Wozniak EA, Chen Z, Paul S, Yang P, Figueroa KP, Friedrich J, et al. Cholecystokinin 1 receptor activation restores normal mTORC1 signaling and is protective to Purkinje cells of SCA mice. Cell reports 2021;37(2).

42. Robinson MD, McCarthy DJ, Smyth GK. edgeR: a Bioconductor package for differential expression analysis of digital gene expression data. bioinformatics. 2010;26(1):139–40.

43. Yu G, Wang L-G, Han Y, He Q-Y. clusterProfiler: an R package for comparing biological themes among gene clusters. Omics: a journal of integrative biology. 2012;16(5):284–7.

44. Subramanian A, Tamayo P, Mootha VK, Mukherjee S, Ebert BL, Gillette MA, et al. Gene set enrichment analysis: a knowledge-based approach for interpreting genome-wide expression profiles. Proceedings of the National Academy of Sciences. 2005;102(43):15545–50.

45. Shami AN, Zheng X, Munyoki SK, Ma Q, Manske GL, Green CD, et al. Single-cell RNA sequencing of human, macaque, and mouse testes uncovers conserved and divergent features of mammalian spermatogenesis. Developmental cell. 2020;54(4):529–47. e12.

46. Green CD, Ma Q, Manske GL, Shami AN, Zheng X, Marini S, et al. A comprehensive roadmap of murine spermatogenesis defined by single-cell RNA-seq. Developmental cell. 2018;46(5):651–67. e10.

