## Supplementary figures and images for "Early gonadotoxic effects of cyclophosphamide on the prepubertal testis and the feasibility of reducing toxicity through combined antioxidant therapy"

### Supplemental Figure 1

#### Supplemental Figure 1.

A

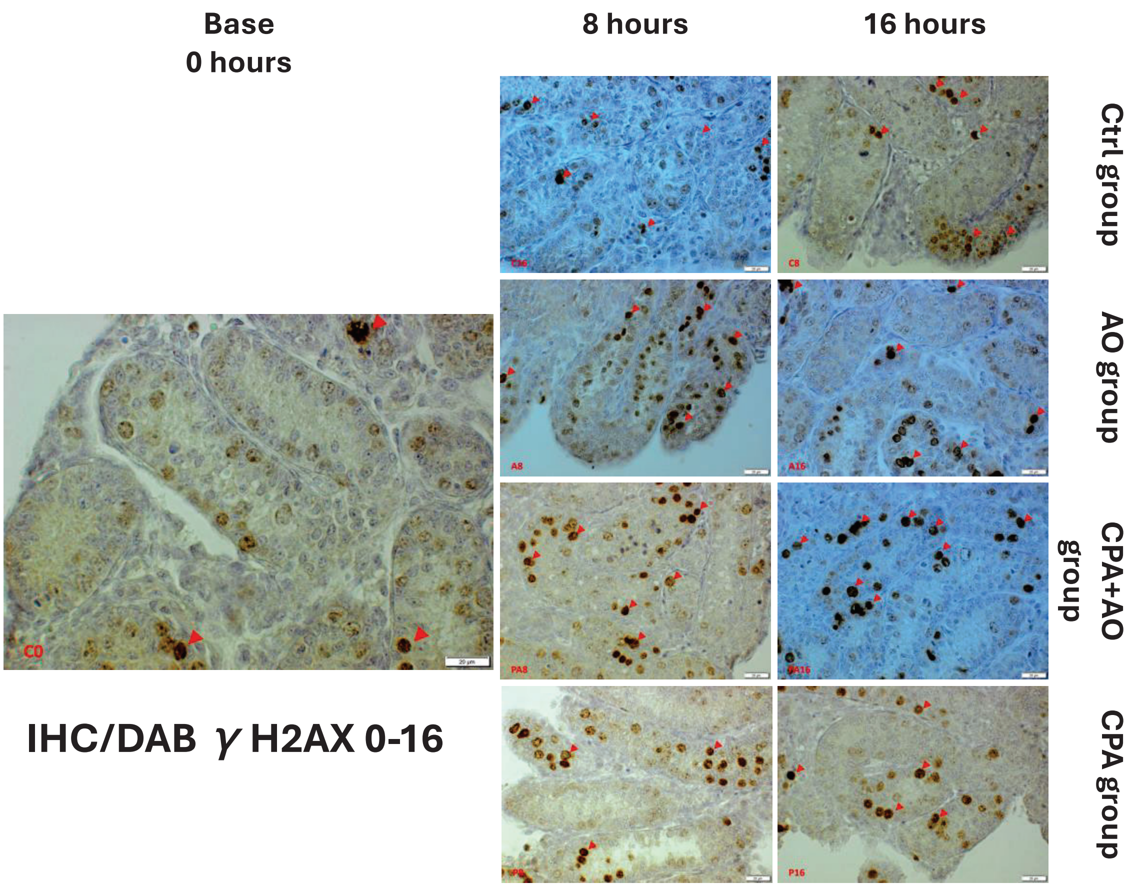


B

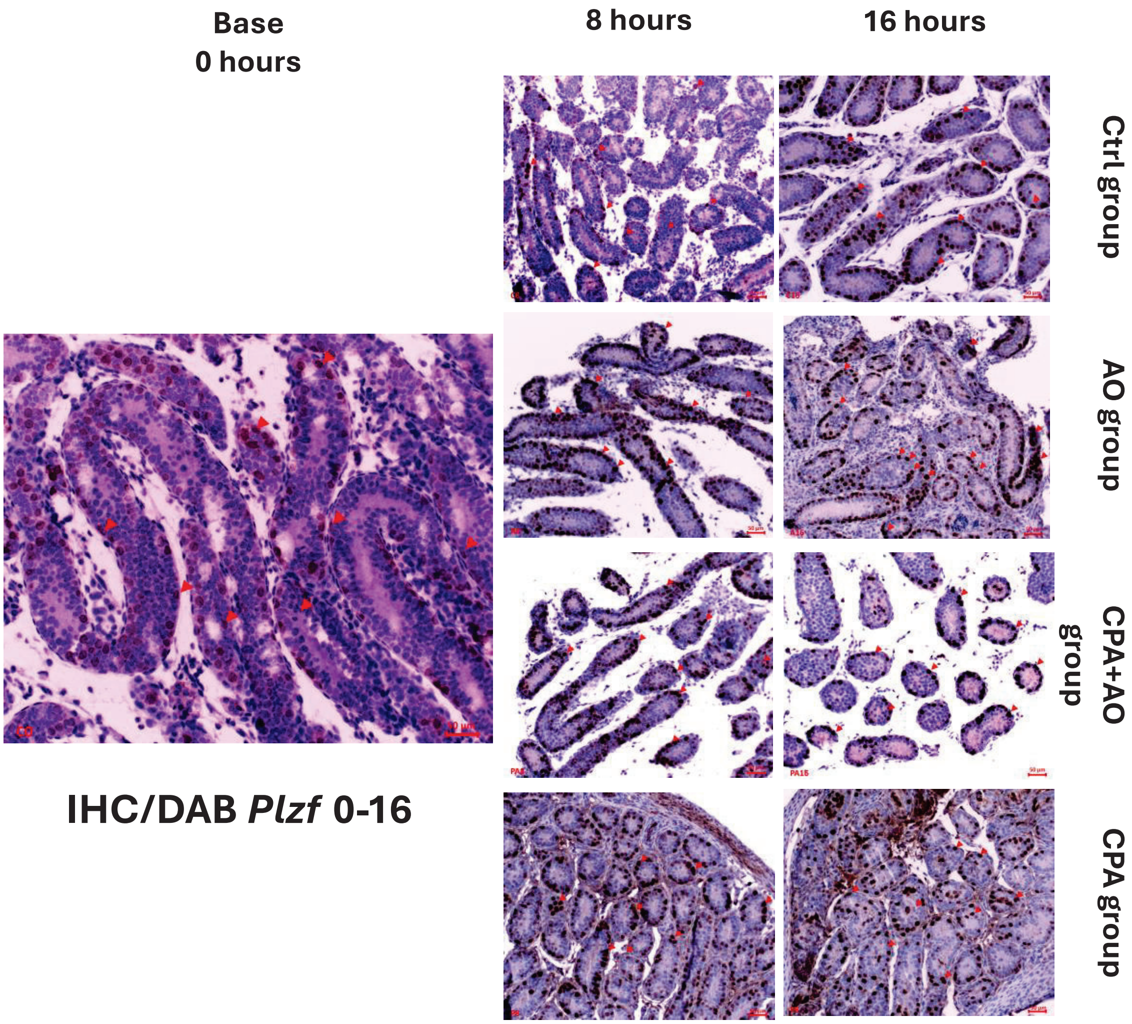

### Supplemental Figure 2

**Supplemental Figure 2.**

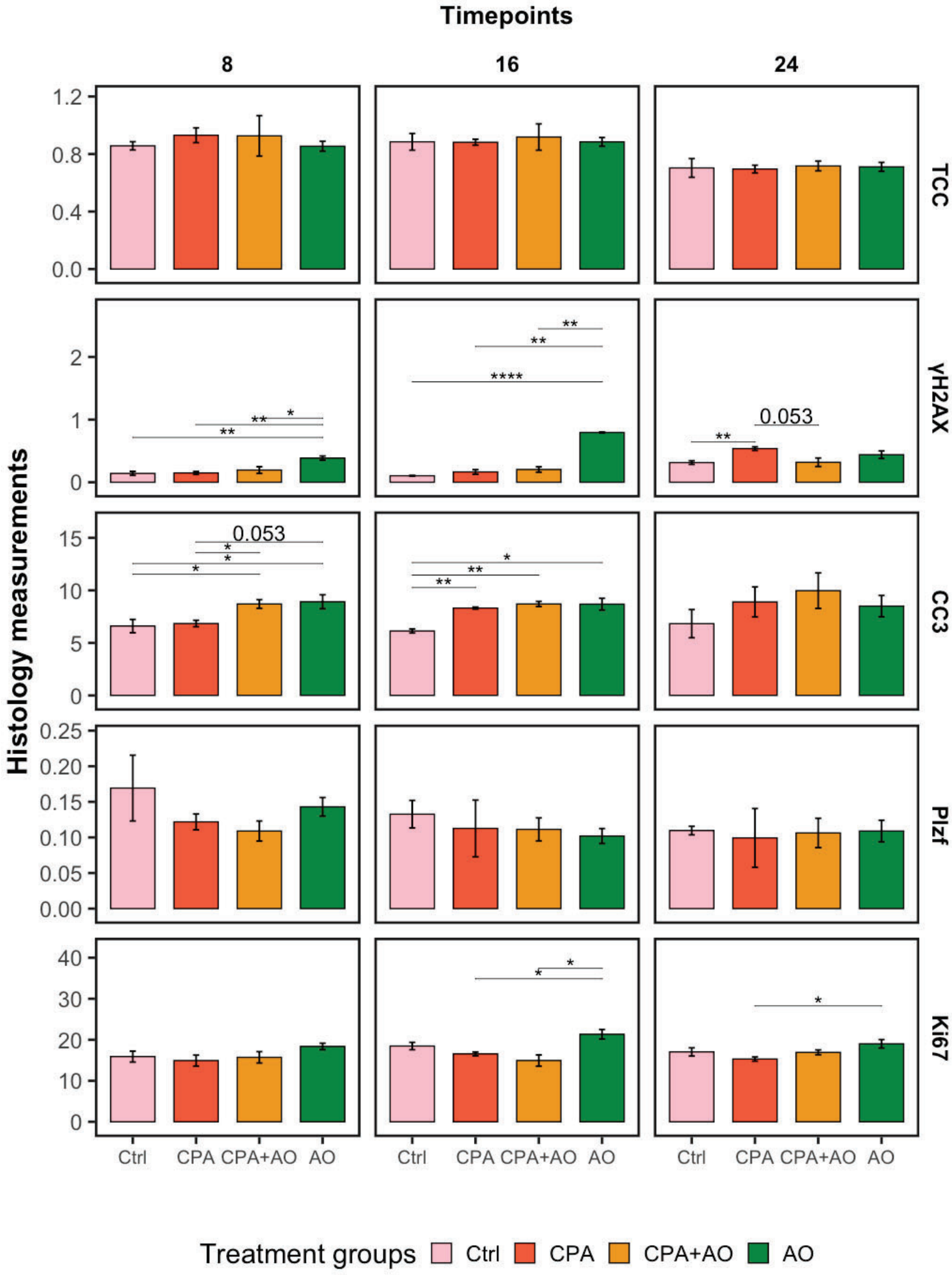
